# Diel predator activity drives a dynamic landscape of fear

**DOI:** 10.1101/221440

**Authors:** Michel T. Kohl, Daniel R. Stahler, Matthew C. Metz, James D. Forester, Matthew J. Kauffman, Nathan Varley, P.J. White, Douglas W. Smith, Daniel R. MacNulty

**Affiliations:** Dept. of Wildland Resources, Utah State University, Logan, UT 84322, USA.; Yellowstone Center for Resources, National Park Service, Yellowstone National Park, WY 82190, USA; Wildlife Biology Program, University of Montana, Missoula, MT 59812, and Yellowstone Center for Resources, National Park Service, Yellowstone National Park, WY 82190, USA; Dept. of Fisheries, Wildlife, and Conservation Biology, University of Minnesota, St. Paul, MN, USA; U.S. Geological Survey, Wyoming Cooperative Fish and Wildlife Research Unit, Dept. of Zoology and Physiology, University of Wyoming, Laramie, WY 82071, USA; Dept. of Biological Sciences, University of Alberta, Edmonton, Alta., Canada T6G 2E9; Dept. of Wildland Resources, Utah State University, Logan, UT 84322, USA

**Keywords:** antipredator behavior, diel activity, elk, habitat selection, landscape of fear (LOF), predation risk, predator activity rhythm, predator-prey interaction, wolf, Yellowstone

## Abstract

A ‘landscape of fear’ (LOF) is a map that describes continuous spatial variation in an animal’s perception of predation risk. The relief on this map reflects, for example, places that an animal avoids to minimize risk. Although the LOF concept is a potential unifying theme in ecology that is often invoked to explain the ecological and conservation significance of fear, quantified examples of a LOF over large spatial scales are lacking as is knowledge about the daily dynamics of a LOF. Despite theory and data to the contrary, investigators often assume, implicitly or explicitly, that a LOF is a static consequence of a predator’s mere presence. We tested the prediction that a LOF in a large-scale, free-living system is a highly-dynamic map with ‘peaks’ and ‘valleys’ that alternate across the diel (24-hour) cycle in response to daily lulls in predator activity. We did so with extensive data from the case study of Yellowstone elk *(Cervus elaphus)* and wolves *(Canis lupus)* that was the original basis for the LOF concept. We quantified the elk LOF, defined here as spatial allocation of time away from risky places and times, across nearly 1000-km^2^ of northern Yellowstone National Park and found that it fluctuated with the crepuscular activity pattern of wolves, enabling elk to use risky places during wolf downtimes. This may help explain evidence that wolf predation risk has no effect on elk stress levels, body condition, pregnancy, or herbivory. The ability of free-living animals to adaptively allocate habitat use across periods of high and low predator activity within the diel cycle is an underappreciated aspect of animal behavior that helps explain why strong antipredator responses may trigger weak ecological effects, and why a LOF may have less conceptual and practical importance than direct killing.

## Introduction

Fear of predation (perceived predation risk) caused by the mere presence of a predator is increasingly regarded as an ecological force that rivals or exceeds that of direct killing (Preisser et al. 2005). The ‘landscape of fear’ (LOF) concept has been advanced as a general mechanism that drives the effects of fear that cascade from individuals to ecosystems (Brown and Kotler 2004, Schmitz 2005, Laundré et al. 2010), including changes in prey physiology (Zanette et al. 2014) and demography (Preisser et al. 2007), plant growth (Ford et al. 2014), and nutrient cycling (Hawlena et al. 2012). Operationally, a LOF is a map that describes the continuous change in predation risk that an animal perceives as it navigates the physical landscape (Brown and Kotler 2004, Laundré et al. 2001, 2010). This mental map of risk overlies the physical terrain like a map of soils, vegetation, or climate, and its ‘peaks’ and ‘valleys’ describe an animal’s perception of those locations as dangerous and safe, respectively (van der Merwe and Brown 2008). Risk perception is indexed by an animal’s measurable response to changes in predation risk (Lima and Steury 2005), and the continuous spatial patterning of this response approximates a LOF as originally defined by Laundré et al. (2001, 2010). Brown and Kotler (2004) defined the concept more narrowly as the spatial distribution of the foraging cost of predation, which is fear measured as the energetic consequence of an animal’s response, chiefly vigilance and (or) time allocation. No matter its definition, the LOF concept is often cited to explain the ecological effects of fear despite two important empirical shortcomings.

First, quantified examples of large-scale LOFs are lacking. Numerous studies have measured animal response to spatial variation in predation risk (reviewed by Moll et al. 2017), but few have mapped this response across physical landscapes as a continuous function of risk in accord with the LOF concept. Among those that have, none mapped areas much larger than 1km^2^ (Shrader et al. 2008, van der Merwe and Brown 2008, Druce et al. 2009, Willems and Hill 2009, Abu Baker and Brown 2010, Emerson et al. 2011, Matassa and Trussell 2011, Iribarren and Kotler 2012, Coleman and Hill 2014). Conversely, some studies have mapped large-scale vegetation patterns and attributed them to animal response to risk without measuring the response itself (Madin et al. 2011). The response has also been overlooked in studies that define a LOF solely in terms of spatial variation in predation risk (e.g., Kauffman et al. 2010, Catano et al. 2016). Large-scale, quantitative examples of a LOF are probably lacking because spatially-explicit data on animal response to risk across vast physical landscapes are difficult to obtain.

Second, little is known about LOF dynamics across the diel (24-hr) cycle. To date, many ecologists have, implicitly or explicitly, assumed that a LOF is a fixed spatial pattern as long as the predator is present (but see Palmer et al. 2017). The underlying rationale is that a constant possibility of predation enforces a chronic state of apprehension in the prey (Schmitz et al. 1997, Brown et al. 1999). This ‘fixed-risk’ assumption of constant attack over time has been a conceptual mainstay in the study of behavioral predator-prey interactions for decades (Lima 2002). Nevertheless, it neglects how predator activity and hunting ability can vary across the diel cycle, and how this may foster a fluctuating acute state of apprehension in the prey and a dynamic LOF despite the constant presence of predators.

Many predators are only active at certain times of day, and visual predators active at night often cannot hunt in absolute darkness. These predatory constraints provide pulses of safety during the diel cycle that may temporarily relieve an animal’s fear of predation and flatten its LOF. This hypothesis is broadly consistent with risk allocation theory, which predicts that animals constantly exposed to predators should respond to pulses of safety with intense feeding efforts (Lima and Bednekoff 1999). It also accords with numerous empirical studies that show how various animals (e.g., zooplankton, rodents, and ungulates) forage in risky places during periods of the diel cycle (e.g., day or night) associated with reduced predator activity and/or hunting ability (reviewed by Lima and Dill 1990, Lima 1998, Brown and Kotler 2004, Caro 2005; see also Fischhoff et al. 2007, Tambling et al. 2012, Burkepile et al. 2013). However, these studies neither tested how animal response to spatial risk is linked to measured variation in diel predator behavior, nor showed how this linkage shapes the animal’s LOF across the diel cycle. Dichotomizing continuous variation in diel predator behavior into periods of presumed safety and danger (e.g., day versus night) is potentially misleading if diel behavior does not conform to these simple categories or if animals assess predation risk as a continuous variable (Creel 2011).

The empirical gaps in the LOF concept are exemplified by its founding case study of elk *(Cervus elaphus)* in northern Yellowstone National Park (YNP) following wolf *(Canis lupus)* reintroduction there in 1995–97 (Laundré et al. 2001). Although this case is frequently cited as a well-understood example of a LOF, and is one that has motivated the proposal that the LOF is a unifying concept in ecology (Laundré et al. 2010), researchers never quantified the elk LOF after wolf reintroduction, nor examined its temporal dynamics in relation to diel wolf behavior. Instead, the elk LOF was inferred from broad-scale, population-level data on vigilance behavior (Laundré et al. 2001), fecal pellets (Hernández and Laundré 2005), and herbivory (Ripple and Beschta 2004) that supported three predictions based on the LOF concept: (1) elk shifted habitat use in response to wolves, including abandonment of high-risk open areas, which (2) decreased diet quality and body fat, and (3) reduced browsing on woody deciduous plants in high risk areas (Laundré et al. 2001, 2010). Some researchers have argued that habitat shifts also reduced elk pregnancy rate (Creel et al. 2009, Christianson and Creel 2014). On the other hand, concurrent fine-scale, individual-level data on movement, body condition, and pregnancy rate indicated elk selected for open areas (Fortin et al. 2005, Mao et al. 2005) and maintained body fat and pregnancy rate (Cook et al. 2004, White et al. 2011, Proffitt et al. 2014). Whereas Fortin et al.’s (2005) 6.5-month study (2001–2002) of 13 female elk equipped with global positioning system (GPS) radio collars suggested elk avoided aspen *(Populus tremuloides)* forests in response to wolves, a three-year experimental study (2004–2007) of aspen demography found that elk browsing was not reduced in risky places (Kauffman et al. 2010). These divergent results have yet to be reconciled, and together they highlight an outstanding need to clarify the elk LOF that prevailed in YNP during the initial years after wolf reintroduction.

The overarching purpose of this study was to improve the empirical foundation of the LOF concept. Our objectives were to (1) quantify a large-scale LOF, and (2) determine how this mental map of risk changes across the diel cycle in response to the daily activity pattern of a predator that is always present. Because the response of Yellowstone elk to wolf reintroduction is a seminal yet unresolved example of a LOF, we examined the elk LOF in northern YNP within the first decade after wolves were released.

We defined the elk LOF as spatial allocation of time away from risky places and times. This conforms to Laundré et al.’s (2001, 2010) broad definition and approximates Brown and Kotler’s (2004) narrower definition. The latter is possible because research indicates that Yellowstone elk manage wolf predation risk mainly through time allocation, keeping vigilance levels constant across habitats that vary in predation risk (e.g., near versus far from forest cover) and increasing vigilance only when wolves are an immediate threat (Childress and Lung 2003; Lung and Childress 2007; Winnie and Creel 2007; Creel et al. 2008; Liley and Creel 2008; Gower et al. 2009; Middleton et al. 2013).

To assess spatial time allocation, we conducted a retrospective habitat selection analysis of data from 27 GPS radio-collared female elk collected during 2001–2004. This included 13 elk from Fortin et al.’s (2005) study, 2 elk from Boyce et al. (2003), 1 elk from Forester et al. (2007, 2009), and 11 elk whose data were never published. Together, these were the first elk GPS location data collected in YNP before or after wolf reintroduction, and we used them to quantify the elk LOF across 995-km^2^ of northern YNP. We tested how this large-scale LOF varied across the diel cycle in relation to the daily activity pattern of wolves which we estimated from direct observations of hunting behavior (1995–2003) and GPS location data (2004–2013). We predicted a dynamic LOF with peaks and valleys that alternated across the diel cycle in response to daily lulls in wolf activity.

## Methods

### Study Area

Our study occurred in a 995-km^2^ area of northern YNP (44° 56’ N, 110° 24’ W) where the climate is characterized by short, cool summers and long, cold winters (Houston 1982). Low elevations (1500–2000 m) in the area create the warmest and driest conditions in YNP, providing important winter range for ungulates, including elk. Vegetation includes montane forest (44%; e.g., lodgepole pine *[Pinus contorta]* and Douglas fir *[Pseudotsuga menziesii*]), open sagebrush–grassland (37%; e.g., Idaho fescue *[Festuca idahoensis]*, blue-bunch wheatgrass *[Pseudoroegneria spicata]*, and big sagebrush *[Artemisia tridentata])*, upland grasslands, wet meadows, and non-vegetated areas (19%) (Despain 1990).

### Study Population

We analyzed habitat selection behavior of 27 adult (> 1 year-old) female elk that spent winter in northern YNP and adjoining areas of the Yellowstone River valley outside YNP from about 15 October to 31 May, 2001–2004. These elk were from a migratory population that numbered from 8,300–13,400 individuals. Our sample of adult female elk was captured in February (2001–2003) via helicopter net-gunning (Hawkins and Powers, Greybull, Wyoming, USA; Leading Edge Aviation, Lewiston, Idaho, USA) and fitted with Telonics (Telonics, Mesa, Arizona, USA) or Advanced Telemetry Systems Inc. (Isanti, Minnesota, USA) GPS radio-collars *(x* ± SD location error = 6.15 ± 5.24 m; Forester et al. 2007) programmed to collect locations at 4–6 hour intervals (5 hour intervals: *n* = 23; alternating between 4 and 6 hour intervals: *n* = 4). To control for movements associated with migratory behavior, we limited our analysis to winter locations collected from 1 November – 30 April. If individuals arrived on the winter range after 1 November, data were censored to the individual’s arrival date (1–22 November). Location data for each individual were collected for 30–353 days *(x* ± SD = 124.5 ± 12.5) across 1–3 winters until collar failure, collar removal, or animal death. We censored location data to include only high-quality locations following guidelines developed by Forester et al. (2009).

Elk age was estimated using cementum analysis of an extracted vestigial tooth (Hamlin et al. 2000) and pregnancy was determined from a serum sample using the pregnancy-specific protein B assay (Sasser et al. 1986, Noyes et al. 1997, White et al. 2011). We evaluated elk nutritional condition via a rump body condition score developed for elk and maximum subcutaneous rump fat thickness measured using an ultrasonograph (Cook et al. 2004). We estimated ingesta-free body fat percentage using the scaled LIVINDEX for elk, which is an arithmetic combination of the rump body condition score and maximum rump fat thickness allometrically scaled using body mass (Cook et al. 2004).

Wolves in this study were members or descendants of a population of 41 radio-collared wolves reintroduced to YNP in 1995–1997 (Bangs and Fritts 1996). The study occurred during a time of peak wolf abundance in YNP: wolf numbers in northern YNP ranged from 70–98 individuals in 4–8 packs (Cubaynes et al. 2014). Each winter, 20–30 wolves, including 30–50% of pups born the previous year, were captured and radio-collared (Smith et al. 2004). Wolves were fitted with very high frequency (VHF; Telonics Inc., Mesa, AZ, USA) or GPS (Televilt, Lindesberg, Sweden; Lotek, Newmarket, ON, Canada) radio-collars. Locations of VHF and GPS-collared wolves were recorded approximately daily during two 30-day periods in early (mid-November to mid-December) and late (March) winter, when wolf packs were intensively monitored from the ground and fixed-wing aircraft, and approximately weekly during the rest of the year. GPS collars recorded locations every hour during the 30-day periods and at variable intervals outside these periods. The proportion of the Yellowstone wolf population that was radio-collared ranged from 35–40%. We captured and handled wolves and elk following protocols in accord with applicable guidelines from the American Society of Mammalogists (Sikes 2016) and approved by the National Park Service Institutional Animal Care and Use Committee.

### Diel activity patterns

We used movement rate to index diel wolf activity given that speed of locomotion is a valid proxy for diel activity patterns in large mammals (Ensing et al. 2014). We estimated movement rate at each hour of the day from the hourly winter positions of 21 GPS-collared wolves recorded in northern YNP during 2004–2013. Wolf GPS data were unavailable prior to 2004. Movement rate equaled the average Euclidean distance of the preceding 1-hour or 5-hour time step. We used hourly movement rate (km/hr) to describe the diel pattern in wolf activity and 5-hour movement rate (km/5-hrs) to test how diel wolf activity influenced elk selection of safe and risky places. We used 5-hour movement rate in the habitat selection analysis to match the 5-hour time interval between consecutive elk locations. To generalize the 1-hour data to 5-hour data, we retained every fifth location beginning with the first 5-hour location available. We used only consecutive 1-hour and 5-hour locations to calculate movement rates.

We estimated the population-level pattern in diel movement rate by applying a generalized additive mixed model (GAMM) to both the 1-hour and 5-hour locations using the mgcv package (version 1.8.0) in R 3.2.3. Because movement data were heavily right skewed (e.g., Fortin et al. 2005), we fit the GAMM using the negative binomial family and incorporated performance iterations such that the scale parameter was as close to 1 as possible. We applied a cyclic cubic regression spline so that the first and last hour of the day matched in accordance with the diel cycle. We included a random intercept for individual identity to account for repeated measures within the study period. We were unable to distinguish between individual and annual variation in wolf diel activity patterns because the number of individuals sampled within years was too small (Appendix S1). Thus, our estimate of diel activity is a population-level estimate calculated as a univariate function of time of day. We used the estimated 5-hour movement rate as the covariate for diel wolf activity in the habitat selection analysis. We used this same approach to model the diel activity pattern of GPS-collared elk, which we did for illustrative purposes. All of our major inferences were based on analyses of elk habitat selection. Each wolf provided an independent measure of movement rate because it was solitary, was the only GPS radio-collared wolf in a pack, or rarely associated with other GPS-collared pack members. The latter was limited to 3 pairs of GPS-collared wolves that were nominally in the same pack during a 30-day period. The proportion of simultaneous fixes that wolves in each pair were near each other (< 2 km) was low: 3%, 6%, and 22%.

We checked that our estimate of diel wolf activity was a valid index of diel hunting pressure during the study period by comparing mean 1-hour diel movement rate to the hourly distribution of daylight (0700–2000) observations of wolves encountering elk in winter from 1995–2003. An encounter was defined as wolves approaching, harassing, chasing, and (or) grabbing elk. Details about how we observed and recorded wolf-elk encounters are described elsewhere (MacNulty et al. 2007).

A concurrent cause-specific mortality study established that wolves were the primary predator of our sample of adult female elk; only one case of cougar-caused mortality was documented (Evans et al. 2006). Analyses of wolf-killed prey during our study period also revealed that elk comprised 90–96% of prey species killed by wolves during winter (Smith et al. 2004; Metz et al. 2012). Together, these studies indicate that the opportunity to kill elk was a key driver of wolf activity in our study area during the period of interest (2001–2004).

### Spatial variation in wolf predation risk

We considered multiple indices of spatial variation in wolf predation risk because it is unclear how elk perceive spatial risk (Beschta and Ripple 2013, Kauffman et al. 2013, Moll et al. 2017). We calculated four indices of spatial risk: predicted occurrence of wolf-killed elk (Kauffman et al. 2007, 2010), density of wolf-killed elk (Gude et al. 2006), openness (Creel et al. 2005, Fortin et al. 2005, Mao et al. 2005), and wolf density (Fortin et al. 2005, Mao et al. 2005, Forester et al. 2007). Kill sites are a well-established metric of predation risk in wildlife systems (e.g., Hopcraft et al. 2005; Thaker et al. 2011; Gervasi et al. 2013; Lone et al. 2014). All spatial risk indices (30 x 30 m grid cell) were developed using the Geospatial Modelling Environment or ArcGIS 10.1.

#### Predicted kill occurrence

We used a previously published model to predict the spatial distribution of wolf-killed elk in northern YNP during each winter of our study (Fig. 1a). Kauffman et al. (2007) developed this model to understand elk response to wolf predation risk in northern YNP. It estimates the relative probability of a kill on the landscape compared to random locations based on the landscape attributes of 774 locations of wolf-killed elk. These kills included all age and sex classes and were documented in winter during a period (1996–2005) that encompassed the present study. Landscape attributes included annual distribution of wolf packs (based on cumulative kernel densities weighted by pack size), relative elk density (from an elk habitat model; Mao et al. 2005), proximity to streams, proximity to roads, habitat openness (forest vs. grassland), slope, and snow depth. The model predicts kill occurrence with respect to the average value of each landscape attribute, such that a predicted kill occurrence of 1 equals no difference between the location of interest and the average landscape, whereas a predicted kill occurrence of 10 equals a kill probability 10 times greater than average for a given year. This produces a year-specific range of values that did not exceed 245 for any year. For example, the range in winter 2000–01 was 0 – 36.5 whereas the range in winter 2001–2002 was 0 – 245.

**Figure 1.**
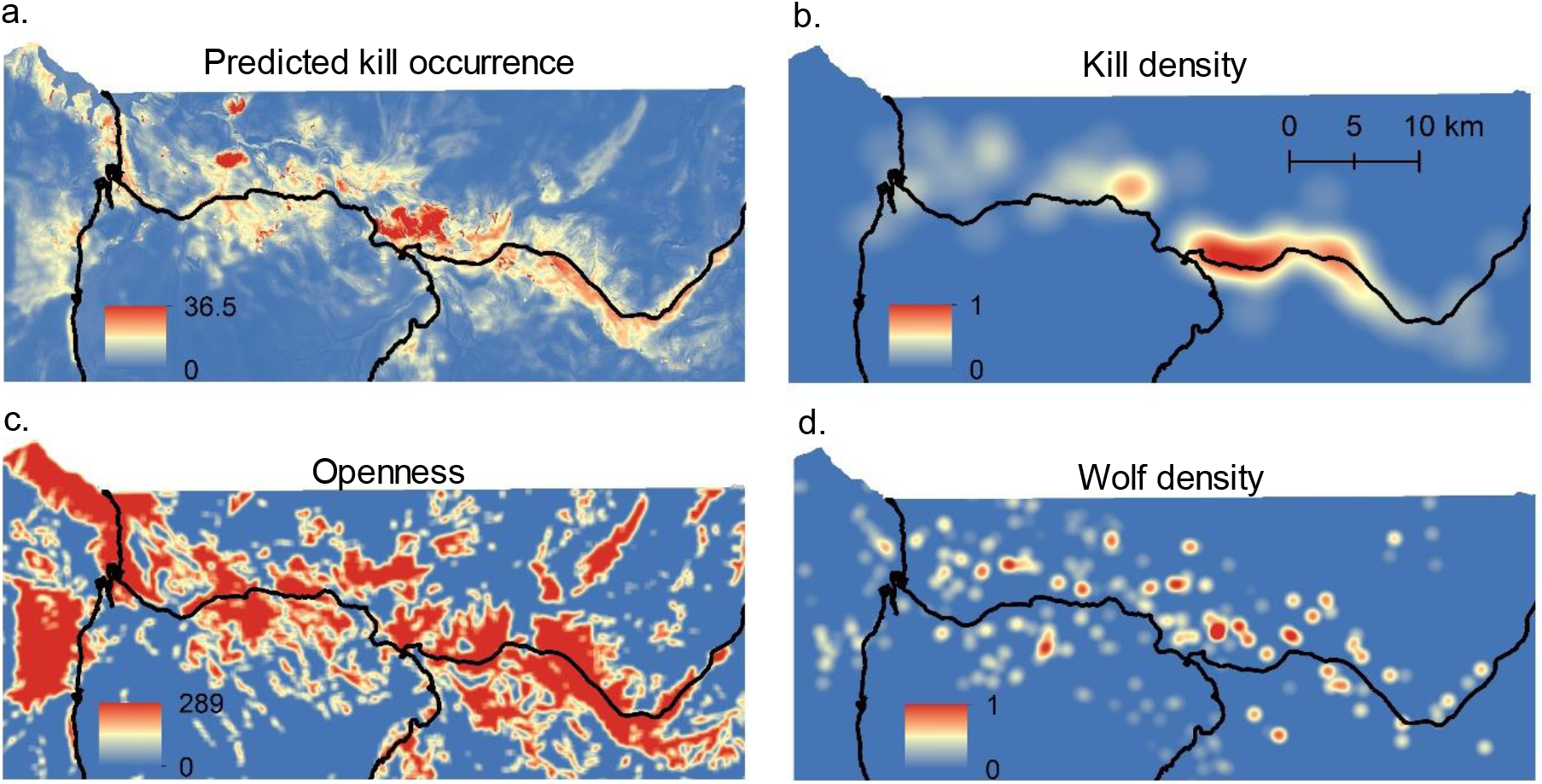
Spatial variation in wolf predation risk during winter in northern Yellowstone National Park was indexed as (a) predicted occurrence of wolf-killed adult male, adult female, and calf elk, (b) density of wolf-killed adult female and calf elk, (c) openness, and (d) density of wolves. (a, b, and d) illustrate conditions during the first year of the study (2001). Openness was consistent across years. Black lines denote roads.

#### Kill density

We used a kernel density estimator (KDE) to estimate the spatial distribution of wolf-killed adult female and calf elk in northern YNP during each winter of our study (Fig. 1b). We excluded kills of adult males because their spatial distribution differed from that of adult females and calves (Pearson’s correlation coefficient, *r* = 0.39; Appendix S2), and we sought to control for possible behavioral responses of adult female elk to sex-specific kill distributions. A total of 235 wolf-killed adult female and calf elk were recorded across the 4 winters (Nov. 2000 – Apr. 2004) following established protocols (Smith et al. 2004). The number of kills included in each annual kill density KDE ranged from 44–84. Following previous studies, we used a fixed bandwidth of 3 km (Fortin et al. 2005). Annual kill density KDEs were standardized from 0 – 1.

#### Openness

We calculated openness (Fig. 1c) as the sum of non-forested cells within a 500 x 500 m moving window centered on each grid cell (range 0 [deep forest] – 289 [open grassland]) following Boyce et al. (2003). We obtained information on the spatial distribution of vegetation types in northern YNP from databases provided by the YNP Spatial Analysis Center. Nonforested pixels were identified from a 1991 vegetation layer which accounted for vegetative changes following the 1988 fires in and near YNP (Mattson et al. 1998). We used this layer to calculate openness because it permitted direct comparison with contemporaneous northern Yellowstone elk habitat selection studies that also utilized the 1991 vegetation layer (e.g., Boyce et al. 2003, Fortin et al. 2005, Mao et al. 2005). We verified that our map of openness was representative of conditions during the study period by comparing it to one calculated from a 2001 LANDFIRE vegetation layer (landfire.gov). We developed and analyzed a single map of openness because there was no inter-annual variation in openness during the study.

#### Wolf density

We estimated wolf density (Fig. 1d) from winter aerial wolf telemetry locations that were randomly filtered to obtain a single location per pack per day. We calculated a least-squares cross-validation fixed smoothing factor (*H*) for each pack with at least 25 locations per winter using Animal Space Use 1.3. Using all non-redundant locations, we used mean *H* (1 km) to calculate annual winter bi-weight kernel densities weighted by pack size (Forester et al. 2007). Annual wolf density KDEs were standardized from 0 – 1.

### Elk habitat selection

We analyzed elk habitat selection using matched case-control logistic regression (CCLR). We used a 1:3 empirical sampling design (Fortin et al. 2005) where, for each end location of a movement step, 3 available locations were sampled with replacement from each individual’s respective step-length and turning-angle distributions. Each set of 4 locations defines a unique stratum (k). Successive strata (k = 10,199) were not independent. Although this autocorrelation does not affect estimated coefficients it does bias the associated standard errors (Fortin et al. 2005). We calculated robust standard errors by specifying an intragroup correlation in our model. Groups were clusters of strata (n = 1,080 clusters) assigned sequentially to each individual each winter and defined by a step-lag at which the autocorrelation was nearly zero. Autocorrelation analysis indicated that this step-lag was 15 steps, such that steps separated by 75 hours were independent (Basille et al. 2015).

We fitted the following CCLR model to all clusters using generalized estimating equations (Craiu et al. 2008):

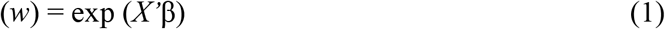

where β is a vector of fitted coefficients and *X* is matrix of explanatory variables for all used and available locations that describe the relative probability of a movement step (*w*), which is the straight-line segment between successive locations at 5-hour intervals. Movement steps with a higher score relative to the set of possible steps have higher odds of being chosen by an animal (Fortin et al. 2005). The sign of the relationship between *w* and spatial risk indicates steps toward or away from risky places: a positive relationship indicates steps toward risky places whereas a negative relationship indicates steps away from risky places. Values of *w* that depict these relationships reflect different levels of perceived predation risk that correspond to the ‘peaks’ and ‘valleys’ in a LOF: minimum values identify peaks (high perceived predation risk) and maximum values identify valleys (low perceived predation risk). We rescaled predicted values of *w* to present an intuitive visualization of the elk LOF (see below).

We could not estimate the main effect of mean 5-hour wolf movement rate because it did not vary within a stratum owing to how used and available locations within a stratum share the same point in time. Within the case-control design of our model, spatial risk variables assigned to each of the three control locations came from the same year in which the use location occurred. Because results did not differ between models fitted to all clusters and models fitted to every other independent cluster *(n* = 2 independent datasets), we present results from the analysis of all the clusters.

For each spatial risk index, we developed a ‘space-only’ habitat selection model and compared it to a ‘space × activity’ model that included terms for the interaction between spatial risk and mean 5-hr wolf movement rate. The space × activity model evaluated how elk selection for risky places at the end of a 5-hour movement step was affected by the mean wolf movement rate during that step. Because prey may not respond instantaneously to predator activity due to imperfect knowledge (Brown et al. 1999), optimal foraging strategies (Kie 1999), shell games (Mitchell and Lima 2002), large landscapes (Middleton et al. 2013a), or a combination thereof, we evaluated the potential for a behavioral lag in habitat selection up to the preceding behavioral step (i.e., 5 hours). We tested different forms of the relationship between habitat selection and spatial risk in the space-only analysis and compared the best-fit space-only model to the best-fit forms in the space × activity analysis. This was necessary to account for how elk in northern YNP may tolerate low levels of spatial risk (Fortin et al. 2005, Mao et al. 2005). We tested for a response threshold by comparing models with a linear effect for spatial risk to models with a threshold effect specified by two linear splines. We performed a grid search of candidate CCLR models to determine the presence and position of thresholds. To control for outliers, we imposed constraints such that the threshold occurred within 1 – 99% of all used data points for a given spatial risk index. This resulted in a range of candidate models (*n* = 41–288) depending on the precision (i.e., decimal units) and scale (i.e., difference in minimum/maximum values) of the spatial risk index.

We compared models using the quasi-likelihood under independence criteria (QIC; Pan 2001), which considers independent clusters of observations while also accounting for nonindependence between subsequent observations (Craiu et al. 2008). The most parsimonious model was the one with the lowest QIC and smallest ΔQIC, which equals the QIC for the model of interest minus the smallest QIC for the set of models being considered. The best-fit model has a ΔQIC of zero.

We performed 1,000 iterations of a 5-fold cross validation for case-control design to evaluate the predictive accuracy of each best-fit model (Boyce et al. 2002). Location data were partitioned into five equal sets and models were fitted to each 80% partition of the data, while the remaining 20% of the data were withheld for model evaluation. Within a cross-validation, the estimated probabilities were binned into 10 equal bins and correlated with the observed proportion of movement steps within the evaluation set. This yielded an average Spearman rank correlation (*r_s_*). Correlations > 0.70 indicate satisfactory fit of models to data (Boyce et al. 2002). CCLR analyses and *k*-folds cross validations were performed in R 3.0.2 using the SURVIVAL and HAB packages, respectively.

### Visualizing the landscape of fear

We used predicted values from our best-fit space × activity step selection model to visualize the LOF for elk in northern YNP. For simplicity, we focused on a single index of spatial risk: kill density. We calculated the predicted relative probability of a movement step (*ŵ*) at each level of kill density at each hour of diel wolf activity. We rescaled these values (1 – *ŵ*) and used the results to elevate the 2-dimensional kill density layer in ArcScene 10.2. Rescaling was necessary so that higher elevations indicated increasing levels of perceived predation risk as per the LOF concept. We constructed a static visualization at two hours when wolf activity was highest (1100: 2.80 km/5-hour) and lowest (1600: 1.42 km/5-hours), and an animated visualization that showed perceived predation risk at each hour of the diel cycle (0000–2300).

## Results

Most GPS-collared wolves (19 of 21) were crepuscular such that their hourly movement rates followed: morning > evening > night > day (Fig. 2a). There was less individual-level variation during peak morning hours than during peak evening hours, indicating that morning was a more reliably active period. The population-average pattern in hourly movement rate during 2004–2013 matched the hourly distribution of directly-observed daylight wolf encounters with elk (r = 0.79; *N* = 502 encounters; Fig. 2a) during 1995–2003. A similar and slightly stronger association was evident when we limited the encounter data to actual kills (r = 0.87, *N =* 89 kills). This suggests that diel variation in wolf movement rate was a meaningful index of diel variation in wolf predation risk. It also suggests, together with evidence that the crepuscular pattern in Fig. 2a was consistent across years (Appendix S3), that the crepuscular pattern during 2004–2013 was representative of the crepuscular pattern during 2001–2004 when elk location data were recorded.

**Figure 2.**
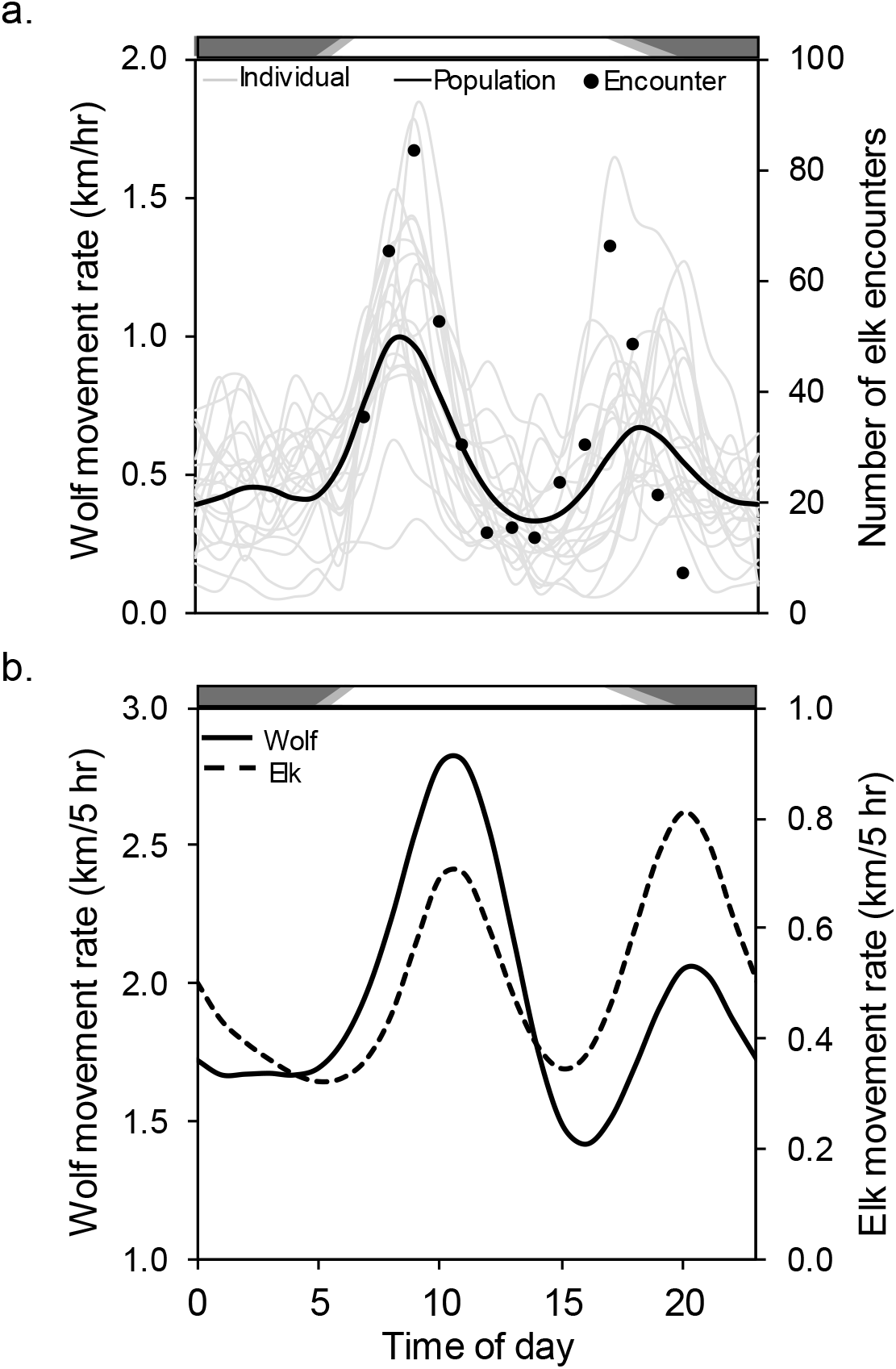
Diel activity patterns of wolves and elk during winter in northern Yellowstone National Park. (a) Mean hourly movement rates for 21 GPS-collared wolves and predicted population mean from a general additive mixed model (left ordinate), and hourly number of directly-observed daylight encounters between wolves and elk (right ordinate). (b) Predicted 5-hr movement rates across 21 GPS-collared wolves (left ordinate) and 27 GPS-collared elk (right ordinate). Bars represent day (white), night (black), and variation in dawn/dusk periods (grey) from 15 Oct. – 31 May.

We estimated wolf movement rate as distance travelled per 5 hours to match the time interval between consecutive elk locations. This shifted the timing of wolf activity to later in the day but it did not alter the crepuscular pattern (Fig. 2b). The mean diel movement rate (km/5-hrs) of elk was similarly crepuscular except that the timing of high and low movement rates was opposite that of wolves: elk movement was greatest at dusk and less at dawn (Fig. 2b). Correlation between wolf and elk movement rates was moderate (r = 0.58).

Irrespective of diel wolf movement, the influence of spatial risk on elk habitat selection was inescapably nonlinear. For each spatial risk index, the best-fit space-only model included a linear spline for spatial risk (Appendix S4), indicating a threshold at which the effect of spatial risk on habitat selection changed. Evidence against a model describing a simple linear relationship between spatial risk and habitat selection was strong for predicted kill occurrence (ΔQIC = 347.13), kill density (ΔQIC = 78.72), openness (ΔQIC = 16.35), and wolf density (ΔQIC = 9.98; Appendix S4). The best-fit models indicated that elk preferred increasingly risky places at low levels of spatial risk *(P* < 0.001; Appendix S5), perhaps due to more food in these areas. At high levels of spatial risk, the effect of risk on habitat selection was negative (wolf density; *P* = 0.02), positive (kill density, *P* < 0.01; openness, *P* < 0.001), or nil (predicted kill occurrence; *P* = 0.76; Appendix S5).

Support for the best-fit space-only models was substantially weaker compared to models that included space × activity interactions between mean diel movement rate (km/5-hrs) of wolves (Fig. 2b) and linear splines for predicted kill occurrence (ΔQIC = 126.73), kill density (ΔQIC = 95.28), openness (ΔQIC = 200.98), and wolf density (ΔQIC = 35.28; Appendix S6). The best-fit space x activity model included a time lag of 2 hour (kill density, openness, wolf density) or 3 hours (predicted kill occurrence; Appendix S6). Five-fold cross validation revealed strong correlations between observed and predicted values for the best-fit space × activity models that included predicted kill occurrence (mean Spearman-rank correlation, *r_s_* = 0.99), openness (rs = 0.99), and kill density (rs = 0.97). Correlations of this magnitude indicate that these models are reliable. By contrast, the reliability of the model that included wolf density was poorer (*r_s_* = 0.67), consistent with earlier findings that wolf density is an inaccurate index of spatial risk in northern YNP due to wolf packs displacing one another from the best hunting grounds where they kill elk (Kauffman et al. 2007). We therefore excluded the wolf density model from further consideration.

Negative space × activity interactions before or after thresholds in predicted kill occurrence (P < 0.001; before threshold), kill density (P < 0.001; after threshold), and openness (P < 0.001; before and after threshold; Appendix S7) showed that elk avoided open grasslands and places where kills occurred when wolf activity was high, but selected for these places when wolf activity was low (Fig. 3a-c). Habitat selection probably did not vary beyond a predicted kill occurrence of 4.5 (Fig. 3a; *P* = 0.87; Appendix S7) because there were few places where the predicted kill occurrence was more than 4.5 times the average kill probability; together, these places comprised only 7% of the study are.

**Figure 3.**
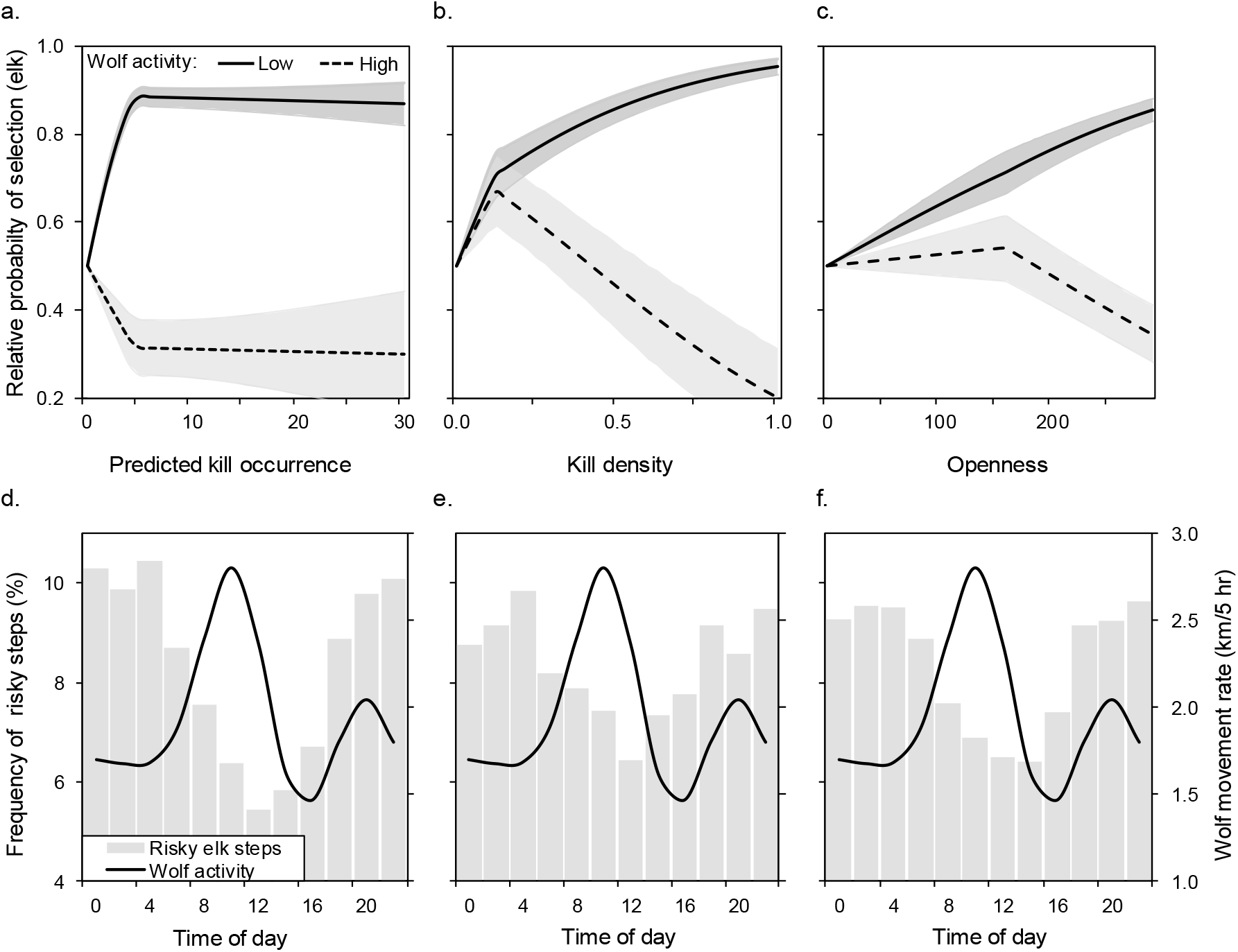
Effects of diel wolf activity (predicted 5-hr wolf movement rate) on elk habitat selection in northern Yellowstone National Park, 2001–2004. (a-c) Elk were more likely to select risky places (areas where kills occurred and open grasslands) when wolf activity was low (1.42 km/5-hrs) than when it was high (2.80 km/5-hrs); lines are population-averaged fitted values with 95% confidence intervals (shaded areas) from the best-fit space × activity models (Appendix S7). (d-f) Frequency of elk steps ending in risky places (locations > mean spatial risk: predicted kill occurrence = 4.5; kill density = 0.22; openness = 194; left ordinate) was greatest at night when wolf activity (mean 5-hr movement rate at 2-hr intervals; right ordinate) was low.

To assess the time of day that elk selected for risky places, we calculated the bi-hourly frequency that elk steps ended in these places. A place was ‘risky’ if it exceeded the average value of a spatial risk index measured across all available locations in the study area. For example, 10.5% of 4084 elk steps ending in places that exceeded the study area’s mean predicted kill occurrence (4.5) happened at 0400–0500, whereas 5.5% of these steps happened at 1200–1300 (Fig. 3d). Steps ending in risky places were most frequent from 2200–0500, which corresponded to the nightly lull in wolf activity (Fig. 3d-f).

To illustrate the effects of diel wolf activity on the elk LOF, we focused on kill density in a portion of our study area (Fig. 4a). Using our best-fit space × activity model for this index (Fig. 4b), we show that places where kills were densely concentrated were valleys (low perceived predation risk) when wolf activity was low (Fig. 4c) and peaks (high perceived predation risk) when wolf activity was high (Fig. 4d). Wolf downtime allowed elk to use places where wolves were more likely to kill them, flattening the LOF every night for about 12 hours (Fig. 3d-f, Video S1). This may explain why prime-aged (2–11 years-old) elk in our sample were in excellent body condition (% ingesta-free body fat; 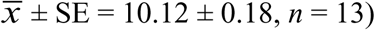 with high pregnancy rates (0.89 ± 0.11, *n* = 15) when radio-collared at midwinter.

**Figure 4.**
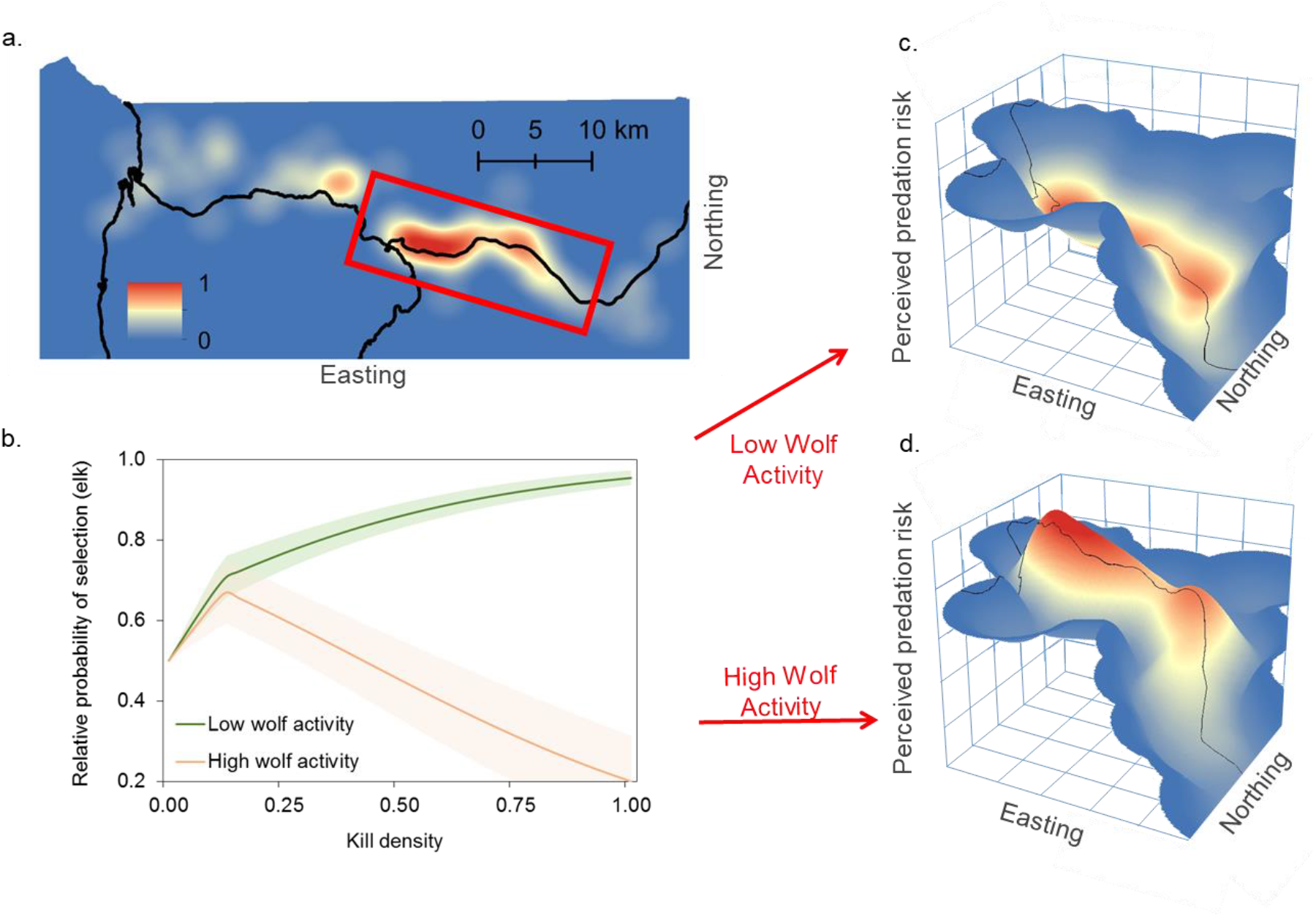
Visualization of how diel wolf activity shaped the landscape of fear for adult female elk in northern Yellowstone National Park, 2001–2004. We examined kill density in one part of our study area, (a), and used the corresponding best-fit space × activity step-selection model, (b), to calculate elk avoidance across this area when wolf activity was low (1.42 km/5-hrs) and high (2.80 km/5-hrs). Risky places where kills were densely concentrated were valleys when wolf activity was low, (c), and peaks when wolf activity was high, (d). Black lines in (a,c, and d) denote roads.

## Discussion

The LOF has been proposed as a possible unifying concept in ecology that explains animal behavior, population dynamics, and trophic interactions across diverse ecosystems (Brown and Kotler 2004, Schmitz 2005, Heithaus et al. 2009, Laundré et al. 2010; Catano et al. 2016). It has also been argued that effective ecological restoration may depend on reestablishing landscapes of fear because fear may be as or more important than direct killing in structuring food webs and modifying ecosystem function (Manning et al. 2009, Suraci et al. 2016). Doubts about the conceptual and practical importance of the LOF stem from a dearth of information about it how it operates across large spatial scales in free-living systems involving apex predators and highly mobile prey (Hammerschlag et al. 2015). We addressed this gap with extensive data from the Yellowstone elk-wolf case study that was the original basis for the LOF concept.

An important aspect of our study is that we measured the LOF as a spatial mapping of time allocation (avoiding risky places and times). This approach accords with the original and widely applied definition of a LOF as a spatial mapping of “any measure of fear” (Laundré et al. 2001, 2010), but differs from the definition of a LOF as a spatial mapping of an animal’s foraging cost of predation (Brown and Kotler 2004). The latter is calculated from giving-up densities which are difficult to measure across vast landscapes like the one we studied (see Bedoya-Perez et al. 2013 for details about the practical uses of giving-up densities). Reconciling the two definitions is important because analyses of a single fear response may describe a landscape that is qualitatively different from a landscape of predation foraging cost, which is an integrative measure of fear that accounts for potential differences in how animal vigilance and time allocation vary with predation risk. For example, if an animal increases its vigilance while foraging in risky places, these places will appear as valleys in a map of time allocation and as peaks in a map of predation foraging cost, thus masking potential ecological effects of fear. Alternatively, if an animal manages risk mainly with time allocation (keeping vigilance constant across safe and risky places), or if vigilance and time allocation respond similarly to temporal variation in risk (decreasing vigilance while foraging in risky places at safe times; Lima and Bednekoff 1999), then the two maps will agree. Constant vigilance provides perfect agreement (Brown 1999), whereas vigilance that covaries with time allocation may provide relatively less relief (lower peaks, shallower valleys) in the map of time allocation, thus underestimating the foraging cost of predation.

Evidence that adult female elk in northern Yellowstone (and adjacent areas) maintain constant vigilance levels across habitats that vary in wolf predation risk (high vs. low wolf densities, near vs. far from forest cover: Childress and Lung 2003; Lung and Childress 2007; Winnie and Creel 2007; Creel et al. 2008; Liley and Creel 2008) suggests our map of time allocation (Fig. 4c-d) matches a map of predation foraging cost. These elk increase vigilance levels only when wolves are an immediate threat (Winnie and Creel 2007; Creel et al. 2008; Lily and Creel 2008; Gower et al. 2009; Middleton et al. 2013) because they can simultaneously process their food and scan their surroundings (Fortin et al. 2004; Gower et al. 2009) as well as escape wolves that attack them (MacNulty et al. 2012; Mech et al. 2015). In general, animals, especially food-limited ones, are expected to use little or no vigilance when they can escape predators in the absence of vigilance (Brown 1999).

On the other hand, if elk vigilance is sensitive to short-term (≤ 24 hours) temporal variation in wolf predation risk as many studies report (Winnie and Creel 2007; Creel et al. 2008; Lily and Creel 2008; Gower et al. 2009; Middleton et al. 2013), then elk may increase vigilance in risky places during periods of the diel cycle when wolves are most active. This is an open question because studies have yet to test how spatial variation in elk vigilance changes across the diel cycle. Nevertheless, theory predicts that an animal’s vigilance level (and its predation foraging cost) should track its predator encounter rate which is itself a function of predator activity level (Houston et al. 1993; Brown 1999; Lima and Bednekoff 1999). If so, elk should reduce vigilance when foraging in risky places during lulls in wolf activity when encounters are infrequent (Fig. 2a) leading to a map of predation foraging cost with more relief than is evident in our map of time allocation (Fig. 4c-d).

We make three important advances with our results. First, we provide a quantified example of a LOF at an unprecedented large scale. Quantified examples are rarer than a casual survey of the literature may suggest because authors often misdefine a LOF as spatial variation in predation risk (e.g., Kauffman et al. 2010, Catano et al. 2016) or an animal’s unmapped response to spatial risk (e.g., Avgar et al. 2015, Hammerschlag et al. 2015, Lyly et al. 2015). Relatively few studies have quantified a spatially-explicit map of an animal’s response to predation risk in accord with the LOF concept. These focused on marine invertebrates (Matassa and Trussell 2011), rodents (van der Merwe and Brown 2008; Abu Baker and Brown 2010), ungulates (Shrader et al. 2008; Druce et al. 2009, Iribarren and Kotler 2012), and primates (Willems and Hill 2009, Emerson et al. 2011, Coleman and Hill 2014) at small spatial scales (< 2 km^2^). Our example is the only one that spans a large-scale (1000-km^2^) landscape. We accomplished this by combining movement data from individually-marked, wide-ranging animals and spatial data describing continuous change in landscape attributes associated with predation risk (kill site locations, vegetation cover). Moving forward, animal-borne transmitters, especially those with accelerometers that permit fine-scale behavioral inferences (Mosser et al. 2014, Collins et al. 2015), together with remotely-sensed spatial risk data (e.g., vegetation cover) may provide the most practical method to estimate landscapes of fear across ecologically-relevant scales.

Second, we demonstrate that diel predator activity is a crucial driver of a LOF. In the large-scale, free-living system we studied, the mere presence of a predator was a necessary but insufficient condition to stimulate a LOF. Had we adopted the classic fixed risk assumption of constant attack over time (Lima 2002) by ignoring diel predator activity, we would have concluded, incorrectly, that our focal prey population had little fear of risky places (Appendix S5). Instead, our consideration of diel predator activity revealed a LOF with peaks and valleys that oscillated across the diel cycle according to the predator’s activity rhythm (Fig. 4, Video S1). This temporally-sensitive response aligns with the ‘risk allocation hypothesis’ (Lima and Bednekoff 1999) which predicts that animals in high-risk environments take maximal advantage of safe times to forage in risky places, and with numerous day-night and light-dark comparisons that show how many taxa (e.g., zooplankton, rodents, and ungulates) use risky places at times of the day when predator activity or hunting ability is minimal (Lima and Dill 1990, Lima 1998, Brown and Kotler 2004, Caro 2005, Fischhoff et al. 2007, Tambling et al. 2012, Burkepile et al. 2013, Palmer et al. 2017).

However, previous studies of diel predator effects on prey habitat use neither quantified a LOF nor linked it to measured variation in diel predator activity as we did. These studies only compared habitat use between day and night, or light and dark periods. This approach would have obscured our results because wolf activity was a complex function of time of day that did not neatly fit the conventional dichotomy of safe and dangerous periods (Fig. 2). As far as we know, our study is the first to quantify how continuous variation in spatial predation risk (Fig. 1) and diel predator activity (Fig. 2) interact with one another to affect an animal’s habitat selection (Appendix S7, Fig. 3) and, ultimately, its LOF (Fig. 4, Video S1). Ecologists have only recently started to investigate the influence of diel predator activity on animal habitat selection (Fischhoff et al. 2007, Tambling et al. 2012, Burkepile et al. 2013). Many of the classic studies of diel predator effects, including zooplankton diel vertical migration (Iwasa 1982) and rodent response to moonlight (Kotler et al. 1991), considered diel changes in the ocular capability of visual predators (Gibson et al. 2009, Upham and Hafner 2013) rather than diel predator activity per se. This aspect of predator-prey interactions deserves more attention because the prevalence of diel activity patterns in apex predators across diverse ecosystems (e.g., Theuerkauf et al. 2003, Roth and Lima 2007, Whitney et al. 2007, Andrews et al. 2009, Cozzi et al. 2012) suggests that it is a potentially common driver of landscapes of fear.

Diel predator activity was an important driver of the landcape of fear in the system we studied because it was a valid source of risk that prey could evidently perceive. Wolves are cursorial hunters that find and select prey by actively searching the environment and visually identifying vulnerable prey that are safe to kill (MacNulty et al. 2007, Mech et al. 2015). As a result, the risk of wolf predation is low when wolves are not highly active. This is illustrated in our data by how the frequency at which wolves encountered, attacked, and killed elk mirrored changes in wolf activity levels (Fig. 2a). The low levels of nightime activity that we documented is consistent with the hypothesis that wolves avoid hunting at night because their vision is best adapted to crepuscular light (Kavanau and Ramos 1975, Roper and Ryan 1977, Theurerkauf 2009). This may explain why wolves in Yellowstone and most other regions exhibit a crepuscular activity pattern (Theurerkauf et al. 2003, Theurerkauf 2009).

The strong statistical association between elk habitat selection and diel wolf activity across three different measures of spatial risk (Appendix S7, Fig. 3) implies that elk perceived diel variation in wolf activity. How elk did this is not obvious from our data. The lagged influence of wolf activity on elk habitat selection (Appendix S7, Fig. 3d-f) suggests that elk did not perfectly perceive changes in wolf activity. Or it could reflect a deliberate tradeoff between safety and food in which elk accepted a higher likelihood of wolf encounter in exchange for more time in preferred foraging habitats. Support for this hypothesis is given by the temporal distribution of elk steps in risky places, which shows that elk minimized their steps in risky places after wolf activity peaked in the morning and started increasing their steps back into these places before wolf activity dipped in the afternoon (Fig. 3d-f). Elk probably tolerate a modest likelihood of wolf encounter because they often survive encounters (MacNulty et al. 2007, Mech et al. 2015). The success of wolves hunting elk in northern YNP during the study period rarely exceeded 20% (Smith et al. 2000, Mech et al. 2001) and dropped below 10% when wolves selected adult elk (MacNulty et al. 2012).

Our third key advance is that we provide the first approximation of the elk LOF that prevailed in northern YNP following wolf reintroduction in 1995–1997. This matters to the discipline of ecology and the practice of conservation because this particular case study is an empirical cornerstone in the LOF concept (Laundré et al. 2001, 2010). Moreover, this case study is a seminal example in the broader debates about the ecological consequences of fear (Ripple and Beschta 2004, Zanette et al. 2011) and the importance of apex predators to the structure and function of ecosystems (Terborgh and Estes 2010, Dobson 2014). Our central finding is that wolves established an elk LOF that was not as relentlessly intimidating as originally proposed and subsequently argued. On the contrary, our results indicate that wolves established a dynamic LOF that shifted hourly with the ebb and flow of wolf activity. Whereas previous studies reported that elk behaviorally abandoned risky places in response to the mere presence of wolves, our research reveals that elk maintained regular use of these areas during nightly lulls in wolf activity. This finding is important because many hypotheses about the ecological effects of the elk LOF in the Greater Yellowstone Ecosystem (GYE) assume that elk abandon risky places when wolves are present.

For example, the ‘predator-sensitive food hypothesis’ that fear of wolves decreases elk pregnancy rate via increased over-winter fat loss assumes that elk move into the protective cover of nutritionally-improverished forests when wolves are present, reducing their use of preferred grassland foraging habitats that have high predation risk (Creel et al. 2009). Although our study is the first to show how elk can safely use grasslands when wolves are present, prior studies of 243 radiocollared elk across four GYE populations (northern Yellowstone, Madison headwaters, Lower Madison, Clarks Fork) have already demonstrated that wolf presence does not prevent elk from using grassland habitats (Fortin et al. 2005, Mao et al. 2005, Proffitt et al. 2009, White et al. 2009a, Middleton et al. 2013a). Evidence that wolves exclude elk from grasslands is limited to a 6.5-month study of 14 GPS-collared elk across two winters (2002–2003) in the Gallatin population (Creel et al. 2005), and a two-month study of elk fecal pellet density across two summers (1998–1999) in northern Yellowstone (Hernandez and Laundré 2005). Decreased elk pellet density with distance from forest edge has been interpreted as evidence that “elk made a significant shift toward the forest edge” following wolf reintroduction (Laundré et al. 2010). This inference is questionable because fecal pellet counts are prone to bias from observer error and variation in fecal disappearance rates (e.g., Campbell et al. 2004, Jenkins and Manly 2008). It also has little bearing on the predator-sensitive food hypothesis which concerns changes in winter habitat use (Creel et al. 2009).

Fortin et al.’s (2005) 7-month study of 13 GPS-collared elk across two winters (2001-2002) in northern YNP is also frequently cited as evidence that wolves exclude elk from grasslands (e.g., Schmitz et al. 2008, Creel et al. 2009, Creel and Christianson 2009, Creel et al. 2011). However, its results are more ambiguous than often acknowledged. Elk were found to prefer conifer forests to grasslands where wolves were numerous, but they were also *more* likely to use grasslands as local wolf densities increased (Fortin et al. 2005: Fig. 3). Confusing matters further, our 26-month study of 27 GPS-collared elk across four winters (2001–2004), which included the 13 animals from Fortin et al. (2005), indicated that wolf density was an unreliable predictor of elk habitat selection (Appendix S6) likely because wolf density was itself an inaccurate gauge of wolf predation risk (Kauffman et al. 2007). These issues highlight the preliminary quality of the results from Fortin et al. (2005).

In winter, our sample of 27 adult female elk used grasslands in northern YNP at night when wolves were relatively inactive (Fig. 3c, 3f). Body fat and blood serum data taken from these elk when radiocollared at mid-winter were consistent with the hypothesis that nocturnal use of preferred grassland foraging habitats was sufficient to offset the effects of wolf presence on elk over-winter fat loss and pregnancy rate. Prime-aged (2–11 yrs-old) animals carried enough body fat (10%) in February to maintain a high rate of pregnancy (89%) contrary to the predator-sensitive food hypothesis. Although our sample is small (<16), the results agree with those from a larger sample of radiocollared elk (>90) from the same population and time period that included the sample we analyzed (Cook et al. 2004; White et al. 2011). They also agree with fetal data from 13,550 adult female, northern Yellowstone elk harvested in Montana (outside YNP) during the 1985–2008 late-season (Dec-Feb) antlerless hunts that indicated pregnancy rate was independent of wolf predation pressure (Proffitt et al. 2014).

Nocturnal use of grasslands may explain how other elk populations utilized these preferred foraging habitats, and why they too maintained relatively high levels of over-winter nutrition and/or pregnancy rate despite wolf presence (Hamlin et al. 2009; White et al. 2009b; Middleton et al. 2013a, b). Counter arguments are based on a potentially unreliable fecal-based pregnancy test of 4 elk populations (Creel et al. 2007, Garrott et al. 2009, White et al. 2011), a snow urine nutritional assay of the Gallatin population over an unspecified time period (Christianson and Creel 2010), and reviews of (un)published data (Creel et al. 2011, 2013). The latter includes a 32% drop in pregnancy rate in the Madison headwaters population (Garrott et al. 2009) that was unrelated to nutrition (White et al. 2009b) and likely an artifact of small sample size and uncontrolled effects of age, which have a profound influence on elk pregnancy rate (Cook et al. 2004, Middleton et al. 2013b, Proffitt et al. 2014). Finally, the consistently crepsucular pattern of wolf activity (Fig. 2, Appendix S3; Theurerkauf 2009) suggests a degree of predictability in wolf predation risk that may explain why wolves have no effect on elk reproduction via chronic stress (Creel et al. 2009, Boonstra 2013).

Elk behavioral abandonment of risky places is also a key mechansism in the behaviorally mediated trophic cascade hypothesis, which asserts that fear of wolves increases productivity of palatable woody deciduous plants in risky places via reductions in elk browsing (Ripple and Beschta 2004, Beyer et al. 2007, Kauffman et al. 2010, Winnie 2012, Peterson et al. 2015). Although population reduction via direct killing could also reduce elk browsing, evidence of an apparent trophic cascade in northern YNP in the decade after wolf reintroduction has been attributed to behavioral mechanisms in part because elk numbers remained high during that period (Ripple et al. 2001, Ripple and Beschta 2004, Ripple and Beschta 2006, Beyer et al. 2007, Ripple and Beschta 2012). We scrutinized the movements of every GPS-collared elk that was tracked in that area during that decade, including 11 previously unreported animals, and our results demonstrate that elk maintained access to aspen *(Populus tremuloides*) and willow *(Salix* spp.) within risky places during daily wolf downtimes. This inference contradicts initial reports from fecal pellet surveys and 13 GPS-collared elk indicating elk avoided aspen where wolves were numerous (Ripple et al. 2001, Fortin et al. 2005). However, it agrees with a winter habitat selection analysis of 80 VHF-collared elk followed in 2000–2002, concurrent to the 13 GPS-collared elk tracked by Fortin et al. (2005), and compared with 94 VHF-collared elk followed before wolf reintroduction in 1985–1990 (Mao et al 2005). This study found that elk *preferred* aspen where wolves were numerous depending on slope and snow levels, and that “elk showed no significant change in selection of aspen, which was highly preferred during winter in both pre- and post-wolf reintroduction periods” (Mao et al. 2005: Table 6). Assessing results from Fortin et al. (2005) and Mao et al. (2005) is difficult, however, because both studies relied on an unreliable index of spatial risk (wolf density; Appendix S6) and an unvalidated GIS layer for aspen.

Nevertheless, elk nocturnal use of areas of high predicted kill occurrence in 2001–2004 (Fig. 3d) accords with separate aspen data taken in 2004–2007 that showed aspen in these same areas did not escape browsing (Kauffman et al. 2010). Similarly, elk avoided riparian areas with willow only during dawn periods (Beyer 2006). This behavior may explain why many willow also did not escape browsing (Bilyeu et al. 2008, Marshall et al. 2013, 2014; but see Beyer et al. 2007). Persistent browsing on aspen and willow was probably also related to how many of these plants existed outside of high-risk areas as defined by our indices of spatial risk (Appendix S8). These results, together with evidence that wolf-caused changes in elk distribution arise from wolves removing individuals rather than elk redistributing themselves (White et al. 2009a, 2010, 2012), support the hypothesis that any indirect effect of wolves on woody deciduous plants is mainly the result of a density-mediated trophic cascade (Creel and Christianson 2009, Kauffman et al. 2010, Winnie 2012, Marshall et al. 2014, Painter et al. 2015).

Although our data are the best available information about the role of wolves in shaping the elk LOF in northern YNP during the first decade of wolf recovery, they are limited in at least three ways. First, the 5-hour interval between consecutive elk locations was coarse and a potential source of bias. This possibility is minimized by the fact that several studies have analyzed subsets of our data and established that the 5-hour interval provides a valid basis for understanding elk movement and habitat selection (Boyce et al. 2003, Fortin et al. 2005, Forester et al. 2007, 2009). Second, our estimated diel wolf activity pattern (Fig. 2) was derived from wolf GPS data collected over a 10-year period (2004–2013) that only partially overlapped our elk study period (2001–2004). This was necessary because GPS data for wolves in YNP were not available until 2004, and the number of wolves equipped with GPS collars each year was small (2–5 animals; Appendix S1). Nevertheless, our estimated diel pattern was most likely representative of the diel pattern during the non-overlapping years because it was: (1) correlated with the time of day that we directly observed wolves encountering (r = 0.79) and killing (r = 0.87) elk during the non-overlapping years (Fig. 2a); (2) consistent across the years in which it was measured (Appendix S3); and (3) similar to diel patterns described for other wolf populations (Eggermann et al. 2009, Theuerkauf et al. 2003, 2009, Vander Vennen et al. 2016).

Although wolves were the primary source of mortality for our study population (Evans et al. 2006), our study, like others before it, ignored the possibility that the elk LOF was shaped by multiple predator species (e.g., wolves and cougars). One reason this may be important is if different predator-specific activity schedules (crespuscular versus nocturnal) create conflicting spatiotemporal patterns of predation risk that require prey to prioritize their response to one predator at the expense of increasing their risk to another. In addition, our analysis did not address the long-term dynamics of the elk LOF. Our results could be an artifact of the potentially unique conditions that prevailed during our study period including a large and growing wolf population, a large but shrinking elk population, and moderate to severe drought conditions. Further research is necessary to determine if and how our estimate of the elk LOF may have changed during the second decade of wolves in northern YNP.

In summary, our major insight is that an animal’s spatially-explicit perception of predation risk (i.e., its ‘landscape of fear’) over a large physical landscape tracks the daily activity pattern of its primary predator, enabling the animal to utilize risky places during predator downtimes, which in turn mitigates the impact of fear on animal resource use, nutritional condition, and reproduction. Our results highlight how a LOF in a large scale, behaviorally-sophisticated system like northern YNP is not a simple, unconditional function of a predator’s mere presence. To assume so may overestimate the threat of predation, underestimate the ability of prey to efficiently manage this threat, and exaggerate the ecological effects of fear. We encourage investigators to recognize the potential for free-living animals to adaptively allocate habitat use across periods of high and low predator activity within the diel cycle. This underappreciated aspect of animal behavior can help explain why strong antipredator responses (e.g., movement, vigilance) may have weak ecological effects, and why these effects may not rival those of direct killing. It also provides a basis for understanding why a LOF may have less relevance to conservation and management than direct killing.

## Acknowledgements

This study was funded by the NSF (DEB–1245373; DEB–0078130), U.S. Geological Survey, National Geographic Society, Natural Sciences and Engineering Research Council of Canada, Alberta Conservation Association, Camp Fire Conservation Fund, Yellowstone Park Foundation, and National Park Service. MTK was supported by a S.J. and Jesse E. Quinney Fellowship from Utah State University. JA Merkle and PJ Mahoney provided helpful statistical advice, and JS Brown, BP Kotler, and one anonymous reviewer provided valuable feedback that improved the manuscript. Any use of trade or product names is for descriptive purposes only and does not imply endorsement by the U.S. Government.

## Supporting Information

**Appendix S1.**
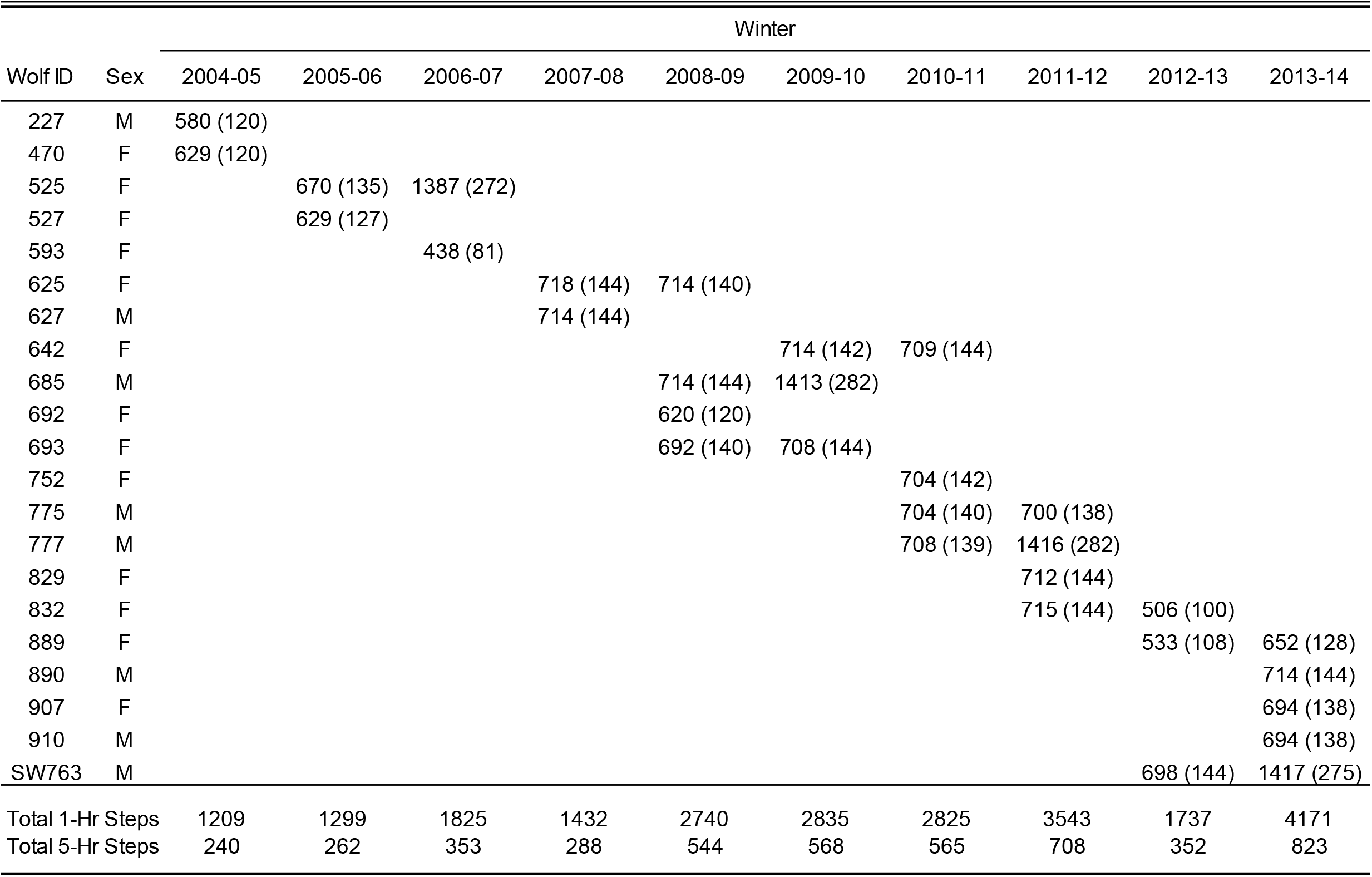
Sample size of movement steps used to calculate diel activity pattern for 21 GPS-collared wolves in northern Yellowstone National Park, 2004–2013. Values represent the steps calculated from consecutive 1-hour (outside parentheses) and 5-hour (inside parentheses) locations.

**Appendix S2.**
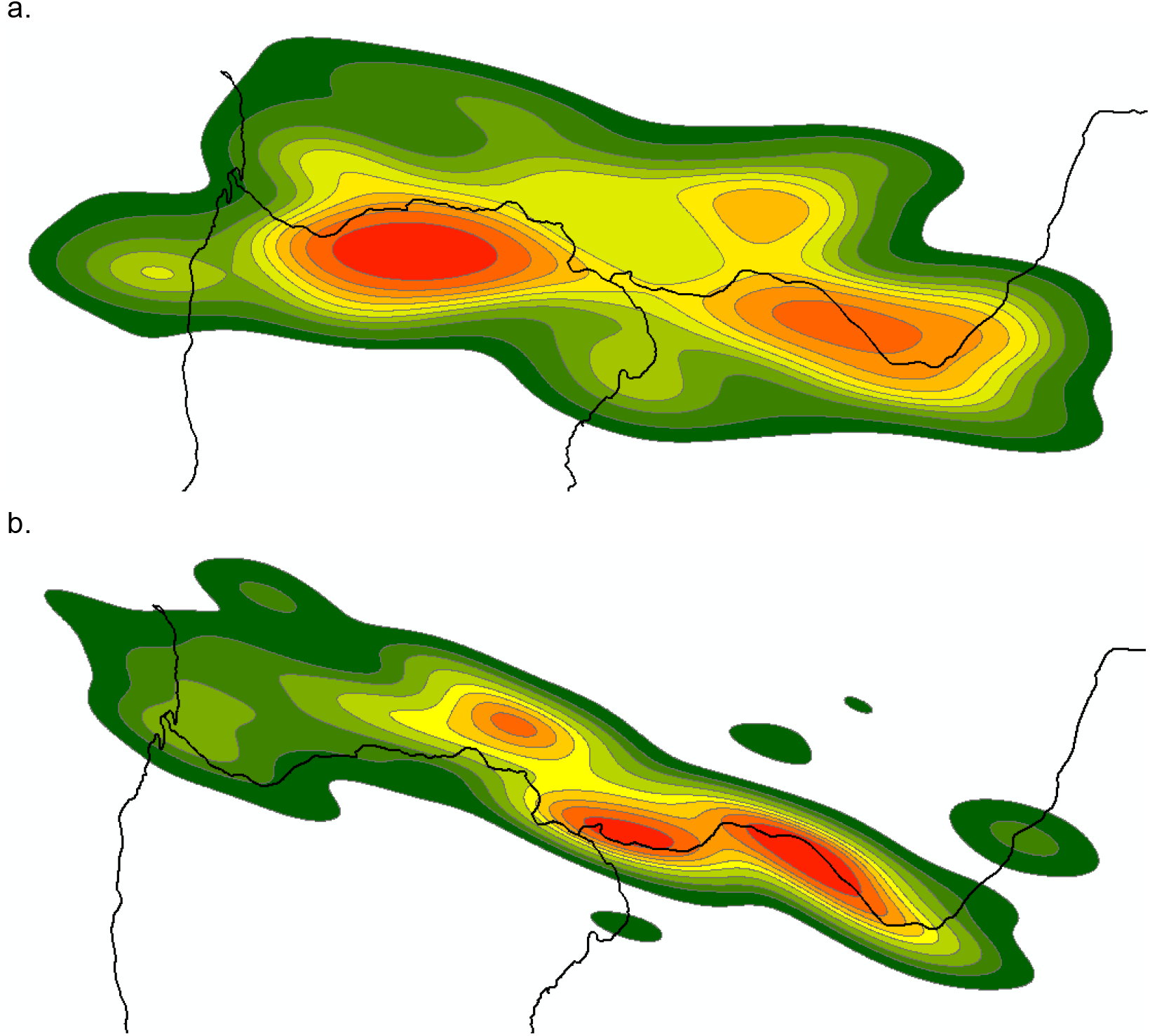
Distribution of wolf-killed (a) adult male elk, and (b) adult female and calf elk during winter in northern Yellowstone National Park, 2001–2004. Contours are 10% kernel isopleths from *a* kernel density estimator applied to kill locations pooled across years. Red rep resents the highest density of kills and black lines denote roads.

**Appendix S3.**
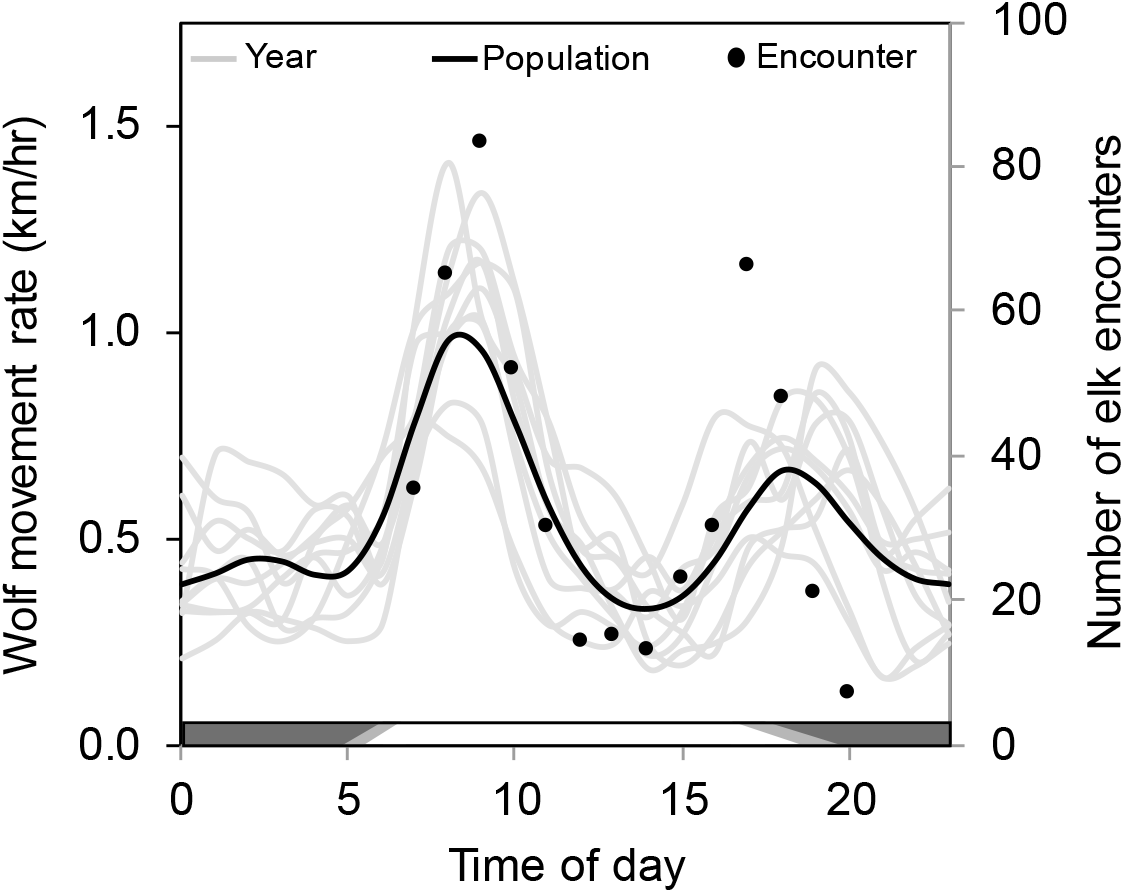
Annual diel activity patterns of wolves during winter in northern Yellowstone National Park, 2004–2013. Mean hourly movement rate for each of 10 years (2–5 GPS-collared wolves per year; Appendix S1) and predicted population mean from a general additive mixed model (left ordinate), and hourly number of directly-observed daylight encounters between wolves and elk (right ordinate). Bars represent day (white), night (black), and variation in dawn/dusk periods (grey) from 15 Oct. – 31 May.

**Appendix S4.**
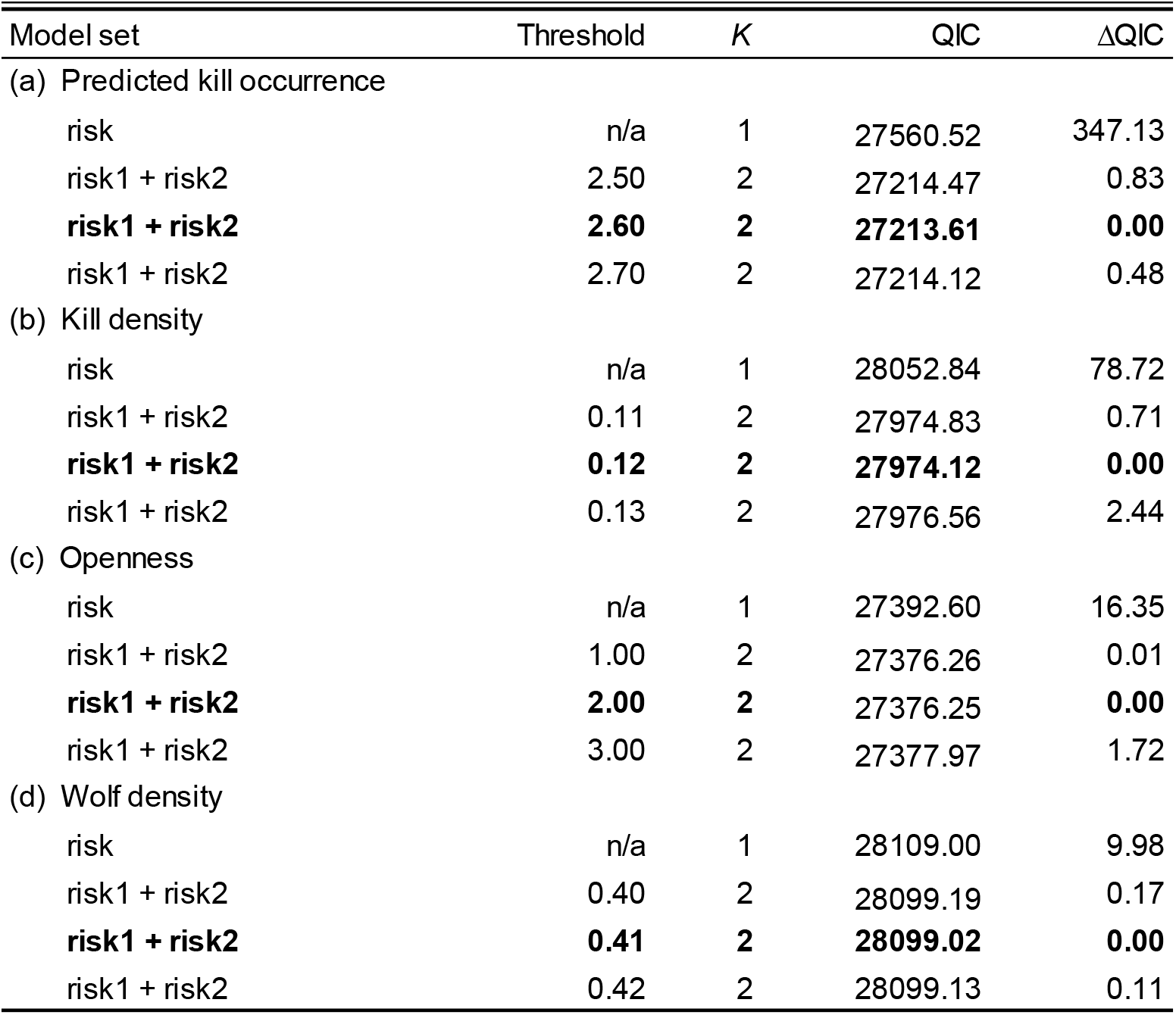
Model selection results for matched case-control logistic regression models describing the relationship between elk habitat selection and four indices of spatial risk (predicted kill occurrence [a], kill density [b], openness [c], and wolf density [d]) in northern Yellowstone National Park, 2001–2004. Variables risk1 and risk2 contain a linear spline for spatial risk at the indicated threshold. The simple linear model (risk) includes no threshold. Number of parameters (K), QIC, and differences in QIC compared to the best model (ΔQIC) are given for each model. The best model for each spatial risk index is in bold face.

**Appendix S5.**
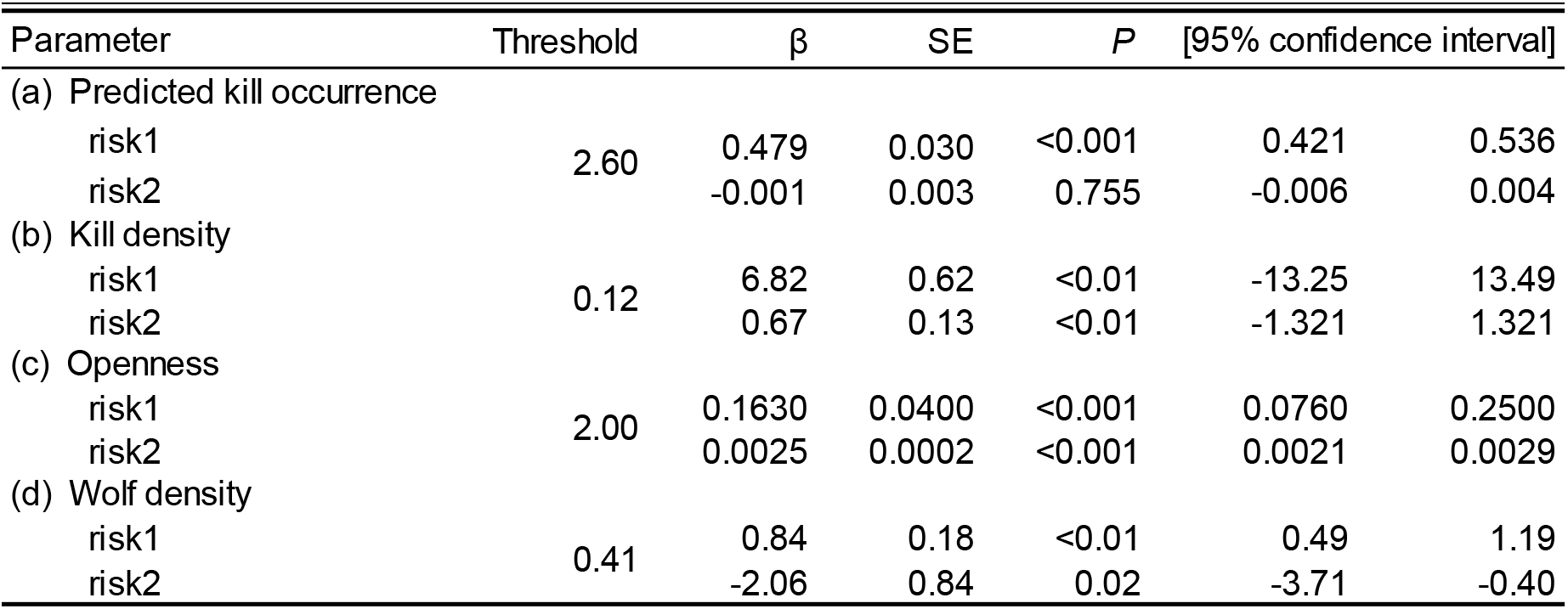
Best-fit matched case-control logistic regression models for the effects of four indices of spatial risk (predicted kill occurrence [a], kill density [b], openness [c], and wolf density [d]) on elk habitat selection in northern Yellowstone National Park, 2001–2004. Variables risk1 and risk2 are the slopes before and after each index-specific threshold. Model selection results are presented in Appendix S4. Confidence intervals were computed using robust standard errors.

**Appendix S6.** Model selection results for matched case-control logistic regression models describing the interactive effect of spatial risk (predicted kill occurrence [a], kill density [b], openness [c], and wolf density [d]) and diel wolf activity (WA; km travelled/5-hr) on elk habitat selection in Yellowstone National Park, 2001–2004. Variables risk1 and risk2 contain a linear spline for spatial risk at the indicated threshold. Space-only models (risk1 + risk2) are the best-fit models from Appendix S5. Space x activity models are the top models from a grid search of thresholds for each hourly lag (≤ 5) in diel wolf activity. Number of parameters (K), QIC, and differences in QIC compared to the best model (ΔQIC) are given for each model. Average Spearman-rank correlation between observed and predicted values calculated from K-fold cross validation (rs) is given for the best-fit model (identified in bold).

| Model set | Threshold | Lag | K | QIC | ΔQIC | $r_s$ |
| --- | --- | --- | --- | --- | --- | --- |
| (a) Predicted kill occurrence |  |  |  |  |  |  |
| risk1 + risk2 | 2.60 | NA | 2 | 27213.61 | 126.73 |  |
| risk1 + risk2 + (risk1 x WA) + (risk2 x WA) | 3.40 | 0 | 4 | 27167.92 | 81.04 |  |
| risk1 + risk2 + (risk1 x WA) + (risk2 x WA) | 4.40 | 1 | 4 | 27126.15 | 39.27 |  |
| risk1 + risk2 + (risk1 x WA) + (risk2 x WA) | 4.50 | 2 | 4 | 27092.01 | 5.13 |  |
| **risk1 + risk2 + (risk1 x WA) + (risk2 x WA)** | **4.50** | **3** | **4** | **27086.88** | **0.00** | **0.99** |
| risk1 + risk2 + (risk1 x WA) + (risk2 x WA) | 4.20 | 4 | 4 | 27117.72 | 30.84 |  |
| risk1 + risk2 + (risk1 x WA) + (risk2 x WA) | 3.40 | 5 | 4 | 27171.30 | 84.42 |  |
| (b) Kill density |  |  |  |  |  |  |
| risk1 + risk2 | 0.12 | NA | 2 | 27974.12 | 95.28 |  |
| risk1 + risk2 + (risk1 x WA) + (risk2 x WA) | 0.12 | 0 | 4 | 27906.66 | 27.82 |  |
| risk1 + risk2 + (risk1 x WA) + (risk2 x WA) | 0.12 | 1 | 4 | 27885.45 | 6.61 |  |
| **risk1 + risk2 + (risk1 x WA) + (risk2 x WA)** | **0.12** | **2** | **4** | **27878.84** | **0.00** | **0.97** |
| risk1 + risk2 + (risk1 x WA) + (risk2 x WA) | 0.12 | 3 | 4 | 27884.87 | 6.03 |  |
| risk1 + risk2 + (risk1 x WA) + (risk2 x WA) | 0.12 | 4 | 4 | 27909.96 | 31.12 |  |
| risk1 + risk2 + (risk1 x WA) + (risk2 x WA) | 0.12 | 5 | 4 | 27948.24 | 69.40 |  |
| (c) Openness |  |  |  |  |  |  |
| risk1 + risk2 | 2.0 | NA | 2 | 27376.25 | 200.98 |  |
| risk1 + risk2 + (risk1 x WA) + (risk2 x WA) | 145.0 | 0 | 4 | 27257.81 | 82.54 |  |
| risk1 + risk2 + (risk1 x WA) + (risk2 x WA) | 147.0 | 1 | 4 | 27208.14 | 32.87 |  |
| **risk1 + risk2 + (risk1 x WA) + (risk2 x WA)** | **159.0** | **2** | **4** | **27175.27** | **0.00** | **0.99** |
| risk1 + risk2 + (risk1 x WA) + (risk2 x WA) | 165.0 | 3 | 4 | 27185.38 | 10.11 |  |
| risk1 + risk2 + (risk1 x WA) + (risk2 x WA) | 2.0 | 4 | 4 | 27242.36 | 67.09 |  |
| risk1 + risk2 + (risk1 x WA) + (risk2 x WA) | 2.0 | 5 | 4 | 27319.06 | 143.79 |  |
| (d) Wolf density |  |  |  |  |  |  |
| risk1 + risk2 | 0.41 | NA | 2 | 28099.02 | 35.28 |  |
| risk1 + risk2 + (risk1 x WA) + (risk2 x WA) | 0.01 | 0 | 4 | 28077.67 | 13.93 |  |
| risk1 + risk2 + (risk1 x WA) + (risk2 x WA) | 0.02 | 1 | 4 | 28069.14 | 5.40 |  |
| **risk1 + risk2 + (risk1 x WA) + (risk2 x WA)** | **0.02** | **2** | **4** | **28063.74** | **0.00** | **0.67** |
| risk1 + risk2 + (risk1 x WA) + (risk2 x WA) | 0.03 | 3 | 4 | 28064.41 | 0.67 |  |
| risk1 + risk2 + (risk1 x WA) + (risk2 x WA) | 0.03 | 4 | 4 | 28071.01 | 7.27 |  |
| risk1 + risk2 + (risk1 x WA) + (risk2 x WA) | 0.03 | 5 | 4 | 28086.39 | 22.65 |  |

**Appendix S7.** Best-fit matched case-control logistic regression models for the interactive effects of spatial risk (predicted kill occurrence [a], kill density [b], and openness [c]) and diel wolf activity (WA; km travelled/5-hr) on elk habitat selection in northern Yellowstone National Park, 2001–2004. Variables risk1 and risk2 are the slopes before and after each index-specific threshold. Model selection results are presented in Appendix S6. Confidence intervals were computed using robust standard errors.

| Parameter | Threshold | Lag | $\beta$ | SE | *P* | [95% confidence interval] | |
| --- | --- | --- | --- | --- | --- | --- | --- |
| (a) Predicted kill occurrence |  |  |  |  |  |  |  |
| risk1 | 4.50 | 3 | 1.103 | 0.072 | <0.001 | 0.96 | 1.24 |
| risk2 |  |  | -0.009 | 0.028 | 0.744 | -0.07 | 0.05 |
| risk1 x WA |  |  | -0.456 | 0.036 | <0.001 | -0.53 | -0.39 |
| risk2 x WA |  |  | 0.002 | 0.015 | 0.873 | -0.03 | 0.03 |
| (b) Kill density |  |  |  |  |  |  |  |
| risk1 | 0.12 | 2 | 8.65 | 3.28 | 0.008 | 2.22 | 15.08 |
| risk2 |  |  | 7.44 | 0.86 | <0.001 | 5.75 | 9.13 |
| risk1 x WA |  |  | -0.95 | 1.65 | 0.564 | -4.18 | 2.28 |
| risk2 x WA |  |  | -3.51 | 0.44 | <0.001 | -4.38 | -2.64 |
| (c) Openness |  |  |  |  |  |  |  |
| risk1 | 159.00 | 2 | 0.011 | 0.002 | <0.001 | 0.006 | 0.015 |
| risk2 |  |  | 0.020 | 0.002 | <0.001 | 0.016 | 0.024 |
| risk1 x WA |  |  | -0.003 | 0.001 | 0.001 | -0.005 | -0.001 |
| risk2 x WA |  |  | -0.009 | 0.001 | <0.001 | -0.011 | -0.007 |

**Appendix S8.**
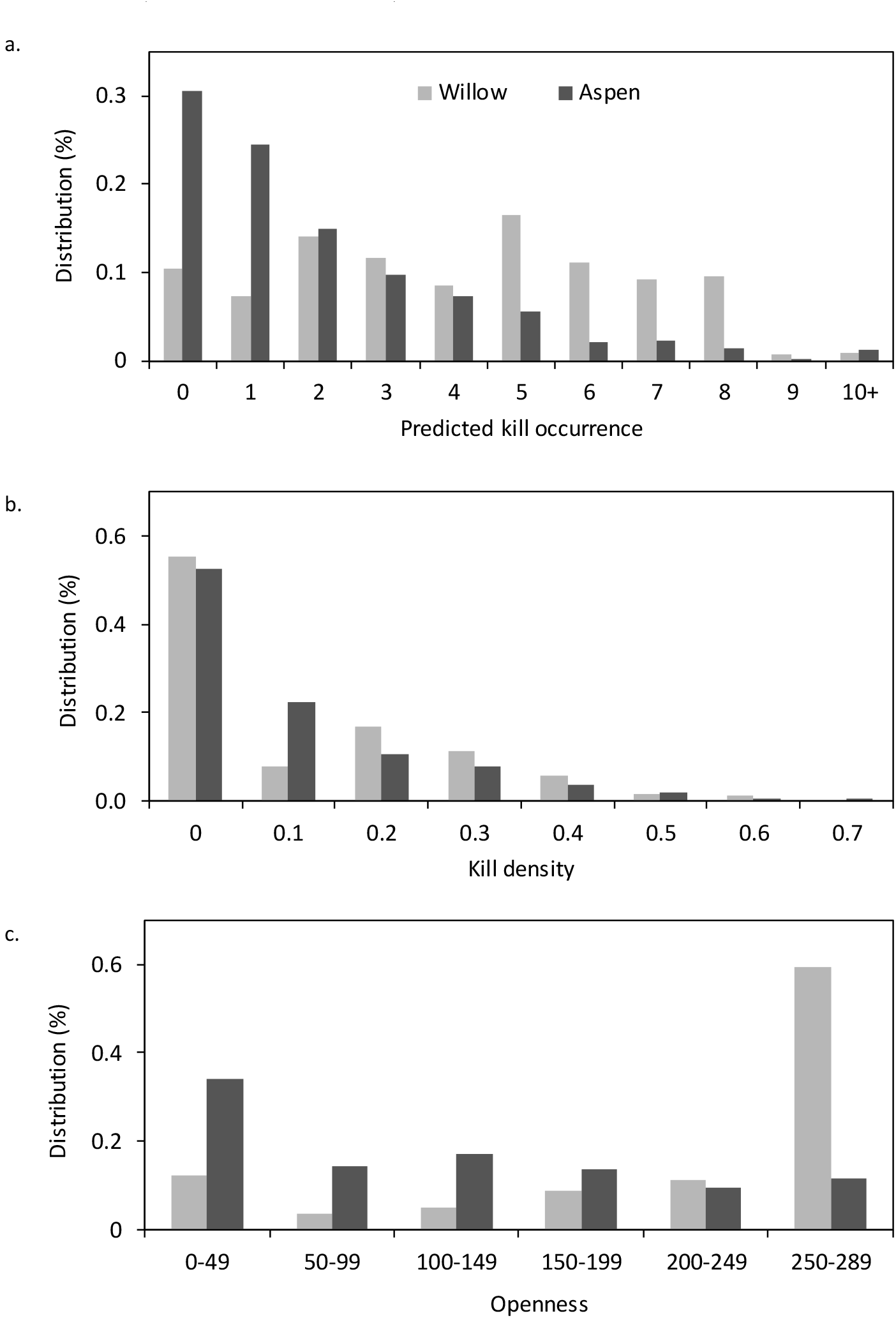
Aspen and willow distribution in northern Yellowstone National Park in relation to spatial variation in wolf predation risk (predicted kill occurrence [a], kill density [b], and openness [c]). Predation risk values in [a] and [b] are the average predicted kill occurrence and kill density at willow and aspen locations from 2000–2004. Aspen location data are from the 1999 Northern Range Vegetation Layer of Yellowstone National Park (Spatial Analysis Center at Yellowstone National Park). Willow location data are from a comprehensive field mapping and inventory that concluded in 2010 (M. Tercek; http://www.yellowstoneecology.com/). Openness data are from a 1991 vegetation layer that accounted for vegetative changes follow the 1988 fires (Mattson et al. 1998).

Video S1. Animated visualization of how diel wolf activity shaped the landscape of fear for adult female elk in northern Yellowstone National Park, 2001–2004. We examined kill density in one part of our study area, (a), and used the corresponding best-fit space × activity habitat selection model, (b), to calculate elk avoidance across this area throughout the diel cycle. Risky places where kills were densely concentrated are represented in red. Peaks identify risky places elk avoided; valleys represent safe places they utilized. Black lines denote roads.

